# Simulating rhizodeposition patterns around growing and exuding root systems

**DOI:** 10.1101/2021.02.25.432851

**Authors:** Magdalena Landl, Adrian Haupenthal, Daniel Leitner, Eva Kroener, Doris Vetterlein, Roland Bol, Harry Vereecken, Jan Vanderborght, Andrea Schnepf

## Abstract

In this study, we developed a novel model approach to compute the spatio-temporal distribution patterns of rhizodeposits around growing root systems in three dimensions. This model approach allows us for the first time to study the evolution of rhizodeposition patterns around complex three-dimensional root systems. Root systems were generated using the root architecture model CPlantBox. The concentration of rhizodeposits at a given location in the soil domain was computed analytically. To simulate the spread of rhizodeposits in the soil, we considered rhizodeposit release from the roots, rhizodeposit diffusion into the soil, rhizodeposit sorption to soil particles, and rhizodeposit degradation by microorganisms. To demonstrate the capabilities of our new model approach, we performed simulations for the two example rhizodeposits mucilage and cit-rate and the example root system *Vicia faba*. The rhizodeposition model was parameterized using values from the literature. Our simulations showed that the rhizosphere soil volume with rhizodeposit concentrations above a defined threshold value (i.e., the rhizodeposit hotspot volume), exhibited a maximum at intermediate root growth rates. Root branching allowed the rhizospheres of individual roots to overlap, resulting in a greater volume of rhizodeposit hotspots. This was particularly important in the case of citrate, where overlap of rhizodeposition zones accounted for more than half of the total rhizodeposit hotspot volumes. Coupling a root architecture model with a rhizodeposition model allowed us to get a better understanding of the influence of root architecture as well as rhizodeposit properties on the evolution of the spatio-temporal distribution patterns of rhizodeposits around growing root systems.

## 2 Introduction

The rhizosphere is defined as the small soil volume around the roots, in which plant roots interact with the soil and thereby alter its physical, chemical and biological properties (Hinsinger et al., 2009). One important rhizosphere process is rhizodeposition, which is defined as the free or passive release of organic compounds by the root, including water-soluble exudates, secretion of insoluble materials and also enzymes such as acid phosphatase, and release of dead root cells (Cheng and Gershenson, 2007). Rhizodeposition affects the ability of plant roots to extract water and nutrients from the soil, which is particularly important when resources are scarce (Hinsinger et al., 2009). Knowledge about the spatial distribution of rhizodeposits in the soil domain is thus crucial (Darrah, 1991).

There are only limited possibilities to directly measure the spatio-temporal distribution patterns of rhizodeposits around a root system. Holz et al. (2018a) used infrared spectroscopy to determine the spatial distribution of mucilage in the rhizosphere. This method allowed them to visualize the axial and radial gradients of mucilage concentration around a single root at a given point in time; information on the temporally dynamic distribution of mucilage is, however, lacking. Under the assumption of a constant ratio between rhizodeposited carbon and root carbon, Pausch et al. (2013) quantified rhizodeposition at the field scale. This approach enabled them to estimate the total amount of rhizodeposition of an entire root system over a defined period of time, however, it does not give any information about the spatial distribution patterns of rhizodeposits.

Simulation models can contribute to better understand the processes leading to rhizodeposition and its spatial and temporal distribution. Such models that describe the distribution of rhizodeposits in the soil domain need to take into account the following processes: the rhizodeposit release by the roots, the diffusion of rhizodeposits into the soil domain, the sorption of rhizodeposits to soil particles and the decomposition of rhizodeposits by microorganisms (Kirk, 1999). A common approach to dynamically compute rhizodeposition patterns in the soil domain is the use of the diffusion-reaction equation. To our knowledge, however, this approach has so far only been applied at the single root scale (Carminati et al., 2016; Holz et al., 2018b; Kirk, 1999) or extrapolated from the single root scale to the root system scale, neglecting differences in rhizodeposition patterns along the root axis (Schnepf et al., 2012). Fletcher et al. (2020) used a citrate-phosphate solubilization model to compute the spatio-temporal distribution of citrate concentrations around root systems in three dimensions. Their approach is, however, limited to very small and simple root systems due to computational limitations.

Various studies have shown the importance of the effect of root architecture on the amount and distribution of rhizodeposits (Hodge et al., 2009; Lynch, 1995; Lynch, Ho, et al., 2005; Manschadi et al., 2014). On the one hand, root architecture controls the amount of rhizodeposit release by the number of root tips (Nielsen et al., 1994). On the other hand, root branching and root growth rate determine whether rhizodeposit release zones can overlap, thereby creating patches of high rhizodeposit concentration, which may facilitate water and nutrient uptake (De Parseval et al., 2017; Holz et al., 2018b).

Rhizodeposition was shown to affect rhizosphere processes such as water and nutrient acquisition only if its concentration exceeds a defined threshold value (i.e., the rhizodeposit hotspot concentration) (Ahmed et al., 2016; Fletcher et al., 2019; Gerke, 2015). However, it is not yet clear when and where around the growing root system such zones of rhizodeposit hotspot concentrations arise, how they are distributed, and what proportion of the total concentration volume they represent. Not only the location of a rhizodeposit hotspot, but also the distance and connectivity to the nearest hotspot and its duration can be a relevant factor controlling soil microbial diversity and microbial activities (Carson et al., 2010). Certain bacteria respond to threats or nutrient availability even when detected from certain distances: volatile organic compounds can provide information over larger distances and diffusible compounds over smaller distances (Schulz-Bohm et al., 2017; Westhoff et al., 2017).

The aim of this study was to couple a root architecture model that simulates the development of a 3D root system with a rhizodeposition model that simulates the transport of rhizodeposits from the root into the soil to investigate the spatio-temporal distribution patterns of rhizodeposits in the soil and to evaluate the influence of root architecture on the generated patterns. For our simulations, we selected the two rhizodeposits citrate and mucilage, which have very distinct properties with regard to the deposition, diffusion, sorption and decomposition rate. In a first scenario, we simulated rhizodeposition by a single growing root. This scenario was used to evaluate the impact of the different rhizodeposit properties such as the rhizodeposit release rate, the sorption to soil particles as well as rhizodeposit decomposition and diffusion on the axial and radial distribution patterns of rhizodeposits around the root. In a second scenario, we investigated the impact of the two root architectural traits ‘root growth rate’ and ‘number of root tips’ on the rhizodeposition patterns around a growing single root and a simple herringbone root system. In a third scenario, we simulated rhizodeposition around the growing root system of *Vicia faba*. This scenario was used to evaluate the impact of a complex root architecture on the spatio-temporal distribution patterns of the rhizodeposits. Additionally, we investigated for how long and where in the soil domain the rhizodeposit concentrations were above a critical threshold value that triggers specific rhizosphere processes, such as an increase in soil water content in the case of mucilage or increased phosphorus mobilization in the case of citrate, and evaluated the importance of root branching and overlap of rhizodeposit release zones for the emergence of such rhizodeposit hotspots. The critical threshold values were thereby selected from literature. In addition, we examined how the distribution of distances from each point in the soil domain to the nearest rhizodeposit hotspot evolves over time.

## 3 Material and Methods

### 3.1 Model development

The simulated root systems consist of root nodes connected by straight root segments, i.e. the explicit 3D root volume is not represented. Roots are therefore considered as point or line sources from which rhizodeposits are released. The possible influence of root diameter on the concentration of rhizodeposits in the soil is thus neglected. In this way, the concentration of rhizodeposits at a given location in the soil domain can be calculated analytically. All equations and assumptions underlying our coupled model approach are explained below.

#### 3.1.1 Root growth model

All root systems were created with the root architecture model CPlantBox, which is described in detail in Schnepf et al. (2018) and Zhou et al. (2020). CPlantBox is a generic model, which allows simulating diverse root architectures of any monocotyledonous and dicotyledonous plant. It distinguishes between different root types, i.e. tap root, basal roots and lateral roots of different order. Each root type is defined by a certain set of parameters that determine its evolution over time. CPlantBox is programmed in C++, but includes a Python binding that allows simplified scripting.

#### 3.1.2 Rhizodeposition model - theory

For each growing root, we solve the diffusion-reaction equation (Jacques et al., 2018) in an infinite domain,

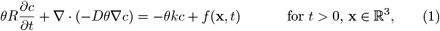

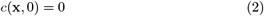

where *θ* is the volumetric water content (*cm*^3^ *cm*^−3^), 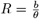 is the retardation factor (*cm*^3^ *cm*^−3^), *b* is the buffer power (−), *c* is the rhizodeposit concentration in the soil (*µg cm*^−3^), *D* = *D*_*l*_*τ* is the effective diffusion coefficient (*cm*^2^ *d*^−1^), *D*_*l*_ is the molecular diffusion coefficient in water (*cm*^2^ *d*^−1^), *τ* is the impedance factor (−), *k* is the linear first order decomposition rate constant (*d*^−1^), *f* is the source term that describes the release of rhizodeposits by the root at position **x** and time *t*.

We consider two cases of rhizodeposition: In the first case, rhizodeposition occurs at the root tip only and the root is thus considered as a moving point source; in the second case, rhizodeposition occurs over a given root length *l* behind the tip and the root is a moving line source. For these two cases, the source term *f* is defined as

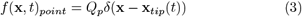

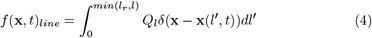

where *Q*_*p*_ (*µg d*^−1^) and *Q*_*l*_ (*µg d*^−1^ *cm*^−1^) are the rhizodeposit release rates of the point and line sources, **x**_*tip*_(*t*) = (*x*_*tip*_, *y*_*tip*_, *z*_*tip*_) is the position of root tip at time *t, l*_*r*_ is the arc length of the exuding root segment (*cm*), **x**(*l*′, *t*) is the position at an arc length of *l*′ behind the position of the root tip at time *t*, and *δ*(**x**) (*cm*^−3^) is the Dirac function.

The analytical solutions to these moving point and moving line source problems have been derived by Carslaw and Jaeger (1959), Bear and Cheng (2010), Wilson and Miller (1978):

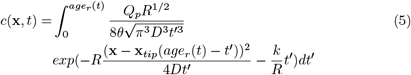

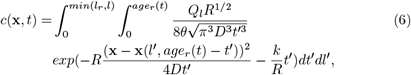

where *age*_*r*_(*t*) is the age of an individual root at time *t* (*d*).

We assume that rhizodeposition stops when the root stops growing. The rhizodeposits, which are already present in the soil, however, continue to diffuse and decompose. Thus, after the root stopped growing, we need to solve:

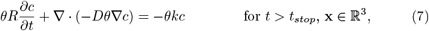

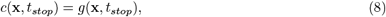

where *g*(**x**, *t*_*stop*_) is the solution concentration (*µg cm*^−3^) at time *t*_*stop*_ (*d*). The analytical solution of the problem with first-order reaction term given by equations (7) and (8) can be derived from the general solution of the homogeneous initial value problem (Evans, 1998) by making use of the transformation *c*′ = *c* ×*exp*(−*k/R* × *t*) (Crank, 1979), where *c*′ is the general solution of the homogeneous problem (Evans, 1998):

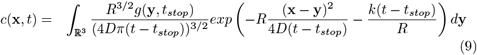

The solution concentration around an entire root system was computed by adding up the concentrations around individual roots, making use of the superposition principle. Thus, the total solution concentration *c*_*T*_ around *N* roots is given by:

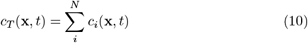

#### 3.1.3 Rhizodeposition model - application

The rhizodeposition model was implemented as an additional module in the root architecture model CPlantBox. The analytical solutions presented in equations (5) and (6) were solved numerically using the Gauss-Legendre quadrature, which we derived from the open source library for C/C++ provided by Pavel Holoborodko (http://www.holoborodko.com/pavel/). This library was used within the C++ code of CPlantBox and introduced into its Python binding so that we could compute the rhizodeposit distribution around a simulated root architecture. The analytical solution for the moving point source (equation (5)) was solved using the function ‘gauss legendre’, while the analytical solution for the moving line source (equation (6)) was solved using the function ‘gauss legendre 2D cube’ with 10 integration points per 1 *cm* root length. The volume integral in equation (9) was solved by trapezoidal rule over a regular cubic grid of 1 *mm* edge length, and the integral was scaled in order to achieve mass balance for diffusion.

To reduce computational time, equations (5) and (6) were not evaluated for the entire soil domain, but only within a specified maximum influence radius around each root within which the rhizodeposit concentrations were significantly different from zero. This maximum influence radius was set to 0.6 *cm* for citrate and to 0.4 *cm* for mucilage, which was a rough estimation of the diffusion length. The rhizodeposit concentrations around an entire root system were computed by adding up the concentrations around individual roots. To reduce computational time, we calculated the rhizodeposit concentrations around the individual roots of the root system in parallel using the multiprocessing package available in Python. In addition, it was necessary to run our model individually for each time step for which an output was needed. We ran all simulations on the Linux cluster of IBG-3 at the Research Center Juelich, which allowed us to run several model runs in parallel. The rhizodeposition model with the code used in this study is publicly available at https://github.com/Plant-Root-Soil-Interactions-Modelling/CPlantBox/tree/publandl2021.

### 3.2 Scenario setup and model parameterization

In a first scenario, we simulated rhizodeposition by a single growing root. This scenario was used to investigate the radial and axial distribution of rhizodeposits around the root. In this scenario, the root was assumed to grow straight downwards at a constant growth rate of 1 *cm d*^−1^ until a root length of 10 *cm* was reached. The root then stopped growing. Rhizodeposition was computed for the two rhizodeposits citrate and mucilage, which have very distinct properties. We used mucilage and citrate rhizodeposit release rates of *Vicia faba*. The rhizodeposit release rate is lower for citrate than for mucilage (Rangel et al., 2010; Zickenrott et al., 2016). The diffusion coefficient and the decomposition rate, in contrast, are higher for citrate than for mucilage (Kirk, 1999; Nguyen et al., 2008; Watt et al., 2006). Furthermore, citrate is known to be sorbed to the soil particles (Oburger et al., 2011), while mucilage that is in contact with free water is not (Sealey et al., 1995). While citrate is exuded from the root apex over a length of approximately 5 *cm* (Pineros et al., 2002), mucilage was shown to be deposited from an area of only a few *mm*^2^ right at the tip of the root (Iijima et al., 2003). All rhizodeposit properties were derived from literature and are presented in Table 1.

**Table 1:**
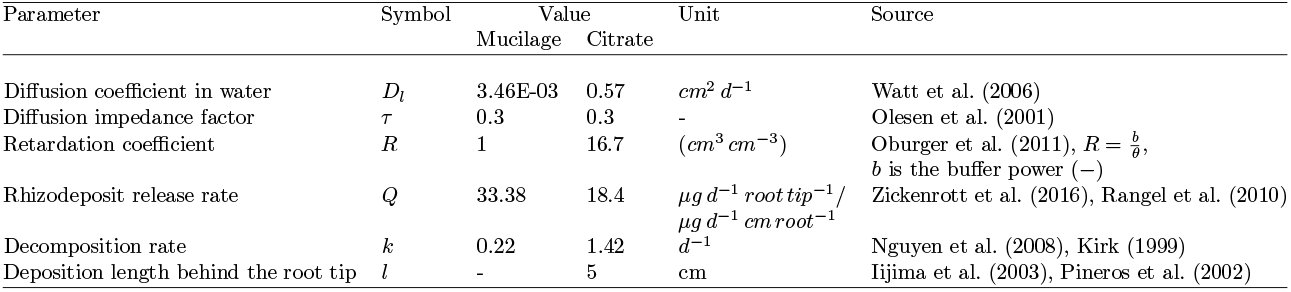
Parameters used in the rhizodeposition model

In a second scenario, we evaluated the impact of the two root architectural traits ‘root growth rate’ and ‘branching density’ on the rhizodeposition patterns around a growing single root respectively a simple herringbone root system. We used four different constant root growth rates (0.1 *cm d*^−1^, 0.5 *cm d*^−1^, 1 *cm d*^−1^, 1.5 *cm d*^−1^) and two different branching densities (2 *cm*^−1^ and 1 *cm*^−1^). Citrate and mucilage rhizodeposit release rates were parameterized for *Vicia faba* using values from the literature (Table 1).

In a third scenario we simulated rhizodeposition by the growing root system *Vicia faba* that was generated with CPlantBox to investigate the impact of a complex root architecture on the spatio-temporal distribution patterns of rhizodeposits. Root architecture parameters were obtained from µCT images of *Vicia faba* plants that were grown in a lab experiment (Gao et al., 2019). The root systems shown on the µCT images were manually reconstructed in a three-dimensional virtual reality system (Stingaciu et al., 2013) and saved as RSML files (Lobet et al., 2015). These RSML files were then used to derive the required input parameters of CPlantBox with the help of a home-grown python code. All input parameters are presented in the supplementary material. The rhizodeposit release rates of citrate and mucilage were adapted to *Vicia faba* using values from the literature and are presented in Table 1. The simulation time was set to 21 days, which is a typical time frame of the lab experiments that were used to image the plant root systems. Simulation outputs were generated in daily time steps. The size of the soil domain was 20 × 20 × 45 *cm*^3^. In all simulation scenarios, the resolution of the soil domain was set to 1 *mm* and we used a constant soil water content of 0.3 *cm*^3^ *cm*^−3^.

To better understand the impact of different plant species on the concentration of rhizodeposits in the soil, we additionally performed simulations for the fibrous root system of *Zea mays* and compared the rhizodeposit mass in the soil domain as well as different root system measures with those of *Vicia faba* in an auxiliary study (see Supplementary Material, Auxiliary study S1).

#### 3.2.1 Rhizodeposit hotspot analysis

Rhizodeposit hotspots are defined as the soil volumes around the root in which the concentration of rhizodeposits is above a critical threshold value and therefore significantly influences specific rhizosphere processes. We defined these threshold values for citrate and mucilage using values from the literature. Gerke (2015) reported that a minimum total carboxylate concentration of 5 *µmol g*^−1^ soil leads to enhanced phosphorus mobilization. Assuming that citrate accounts for about 25 % of the total carboxylate concentration (Lyu et al., 2016) and using the soil buffer power as the ratio between the total rhizodeposit concentration and the soil solution rhizodeposit concentration (Nye, 1966), this corresponds to a threshold citrate concentration of 58 *µg cm*^−3^ soil solution at an assumed bulk density of 1.2 *g cm*^−3^. In a modelling study based on experimental measurements, Carminati et al. (2016) investigated the effect of mucilage on rhizosphere hydraulic properties and transpiration as a function of mucilage concentration. For a sandy soil, they observed a measurable effect of mucilage on soil water retention at a minimum mucilage concentration of 0.33 *mg g*^−1^ dry soil, which corresponds to a threshold mucilage concentration of 1300 *µg cm*^−3^ soil solution at an assumed bulk density of 1.2 *g cm*^−3^. It was shown that not only fresh mucilage, but also mucilage derivatives that are produced during the process of decomposition can have an impact on soil hydraulic properties (Carminati and Vetterlein, 2013; Or et al., 2007). To date, however, it is not clear how mucilage derivatives affect soil water dynamics (Benard et al., 2019). In this study, degraded mucilage is neglected and only the concentration of fresh mucilage is taken into account.

To compare hotspot volumes of root systems that differ in architecture or age, we normalized them with the root length and with the minimum soil volume that contains 99 % of the total rhizodeposit mass that is currently present in the soil domain. These relative hotspot volumes are further on called length-normalized and volume-normalized rhizodeposit hotspot volumes. While the length-normalized hotspot volume is a measure of the efficiency of the root architecture, the volume-normalized rhizodeposit hotspot volume can be regarded as a measure of the efficiency of rhizodeposition.

The duration of an individual rhizodeposit hotspot at a specific location in the soil domain is not constant, but varies depending on different dynamic processes such as the diffusion and decomposition rate, the sorption to soil particles, the deposition length behind the root tip and the root architecture, which may cause rhizodeposit overlap. We therefore also investigated the lifetime of rhizodeposit hotspots within the soil domain.

To examine how the distribution of distances from each point in the soil domain to the nearest rhizodeposit hotspot evolves over time, we applied the 3D ImageJ Suite (Ollion et al., 2013) plugin of Fiji (Schindelin et al., 2012) to calculate the Euclidean 3D distance maps from the nearest hotpots at various days of root growth and provide the histograms of the distance maps.

## 4 Results

### 4.1 Scenario I: Rhizodeposition by a single growing root

Fig. 1 shows the concentration profiles of citrate and mucilage around a growing and exuding single root after a defined time period. After 10 days, the root reached its maximum length of 10 *cm* and both root growth and rhizodeposition stopped. Diffusion and decomposition of the rhizodeposits continued until the end of the simulation. For both citrate and mucilage, the concentrations were thus much higher after 10 days (Fig.1 (I)) than after 15 days (Fig.1 (II)) of simulation due to the ongoing decomposition of the rhizodeposits. The progressive diffusion furthermore led to a larger extent of the radial profiles after 15 days compared to 10 days and also at position 2 (15 *cm* behind the root tip) compared to position 1 (1.5 *cm* behind the root tip). In general, concentrations of mucilage were higher than concentrations of citrate due to the differences in rhizodeposit properties. The peak concentration of mucilage was located at a distance of 1 *cm* behind the root tip, while citrate concentrations were highest 5 *cm* behind the root tip. The radial extension of the concentration from the root axis was larger for citrate than for mucilage due to the larger ratio of the effective diffusion coefficient and the retardation factor (Fig.1 (b,c)). The rhizodeposit hotspot concentrations extended over a length of 5.3 *cm* and 2.2 *cm* along the root axis for citrate and mucilage, respectively, while the root was still growing (Fig.1 Ia). The maximum radial extent of the rhizodeposit hotspot concentration was 1 *mm* and 0.5 *mm* for citrate and mucilage, respectively (Fig.1 Ib, c). The maximum radial extent of citrate and mucilage rhizopheres in which the rhizodeposit concentration was below the threshold value, but still detectable, was 4 − 9 *mm* and 2 − 5 *mm* for citrate and mucilage, respectively (Fig.1 Ib, c).

**Figure 1:**
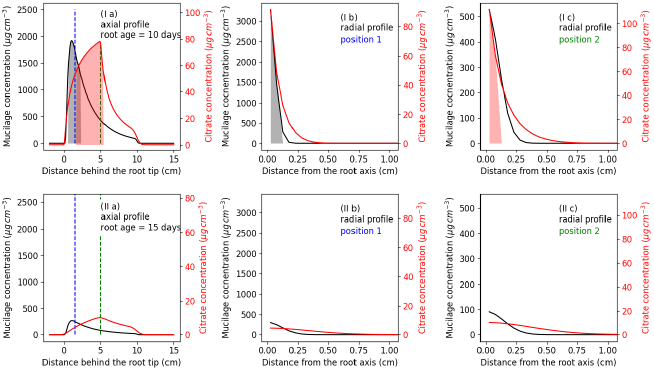
Concentration profiles of mucilage and citrate after (I) 10 and (II) 15 days: along the root axis (a) and radially from the root axis at a distance of 1.5 *cm* (position 1) (b) and 15 *cm* (position 2) (c) from the root tip; the dotted lines specify the location on the axial profile (a) where the radial profiles (b) and (c) were taken; the shaded areas denote the part of the profiles where the concentrations are above the threshold values

### 4.2 Scenario II: Impact of root architectural traits on the rhizodeposition patterns around a single growing root

#### 4.2.1 Impact of root growth rate

Considering that rhizodeposits are released from the growing tip in the case of mucilage and from a small zone behind the growing tip in the case of citrate, changes in root elongation rate had a strong impact on the distribution of rhizodeposits in the soil. In figures 2 and 3 the concentrations of mucilage and citrate around a single straight root that elongates for 10 days at different constant growth rates are shown. A larger growth rate led to a larger soil volume containing rhizodeposits at a lower concentration. In black, we depicted the volume of rhizodeposit hotspots for both citrate and mucilage. The largest rhizodeposit hotspot volume was found for the second lowest root growth rate of 0.5 *cm d*^−1^ for citrate and for the second highest root growth rate of 1 *cm d*^−1^ for mucilage.

**Figure 2:**
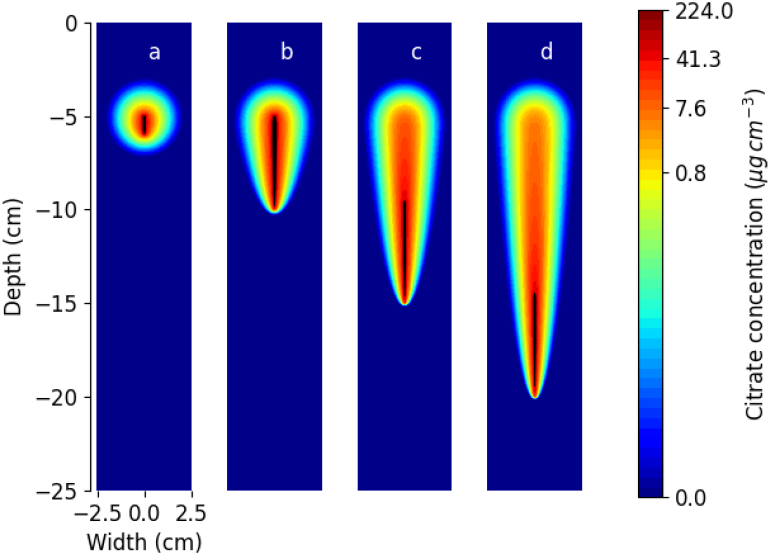
Concentration of citrate deposits around a single root after 10 days of growth at a constant growth rate of (a) 0.1 *cm d*^−1^, (b) 0.5 *cm d*^−1^, (c) 1 *cm d*^−1^, (d) 1.5 *cm d*^−1^. The black patches denote the hotspot volume; note that the colors are in logarithmic scale

**Figure 3:**
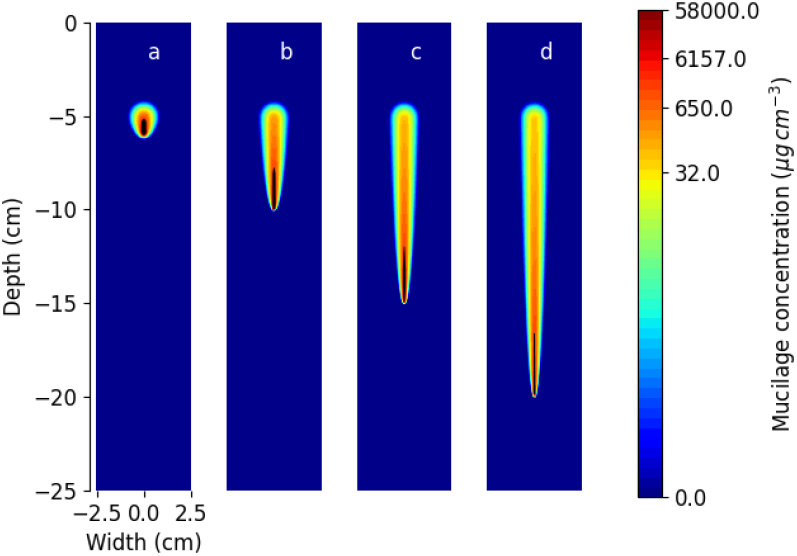
Concentration of mucilage deposits around a single root after 10 days of growth at a constant growth rate of (a) 0.1 *cm d*^−1^, (b) 0.5 *cm d*^−1^, (c) 1 *cm d*^−1^, (d) 1.5 *cm d*^−1^. The black patches denote the hotspot volume; note that the colors are in logarithmic scale

#### 4.2.2 Impact of root branching patterns

Fig. 4 shows the distribution of rhizodeposits around two simple herringbone root systems with different branching densities for both citrate and mucilage. An increase in branching density by a factor of two (from 9 to 16 root tips) increased the total mass of rhizodeposits present in the soil domain by 48 % for citrate and by 79 % for mucilage after 10 days of growth. There were no rhziodeposit hotspot volumes (depicted in pink) around the upper laterals and the citrate rhizodeposit hotspot volumes were located further behind the root apex than the mucilage rhizodeposit hotspot volumes. An increase in branching density by a factor of two increased the total rhizodeposit hotspot volume by 80 % and 73 %, the length-normalized hotspot volume by 13 % and 9 % and the volume-normalized hotspot volume by 51 % and 29 % for citrate and mucilage, respectively, after 10 days of growth. For our parameterization, root branching thus had a greater impact on the total rhizodeposit hotspot volume and also on the rhizodeposition efficiency of citrate than of mucilage.

**Figure 4:**
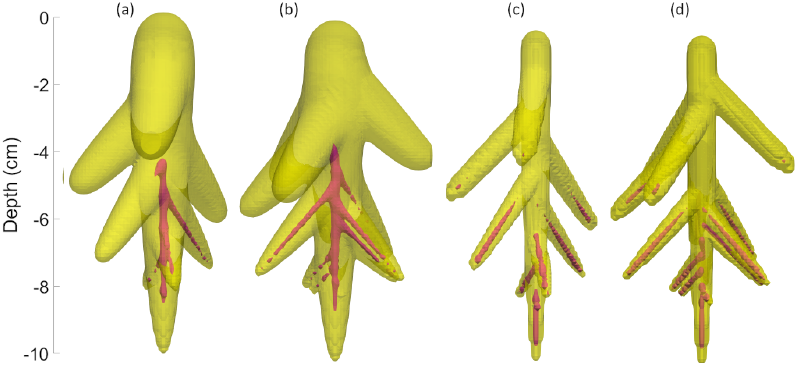
Deposition patterns of rhizodeposit hotspot concentrations (pink) and concentrations above the arbitrary threshold of 0.1 *µg cm*^−3^ (yellow) for citrate (a,b) and mucilage (c,d) around a simple herringbone root system with different branching densities (1 *cm*^−1^ (a,c) and 2 *cm*^−1^ (b,d)) after 10 days of growth at a constant growth rate of 1 *cm d*^−1^

### 4.3 Scenario III: Rhizodeposit concentration patterns around the root system of *Vicia faba*

Fig. 5 shows the rhizodeposit concentration patterns of citrate and mucilage around the 21 day old root system of *Vicia faba*. The maximum extent of the rhizosphere was defined using an arbitrary threshold of 0.1 *µg cm*^−3^. The maximum mucilage concentrations were larger than the maximum citrate concentrations and the extent of the citrate rhizosphere (Fig. 5 (a)) was larger than the extent of the mucilage rhizosphere (Fig. 5 (b)).

**Figure 5:**
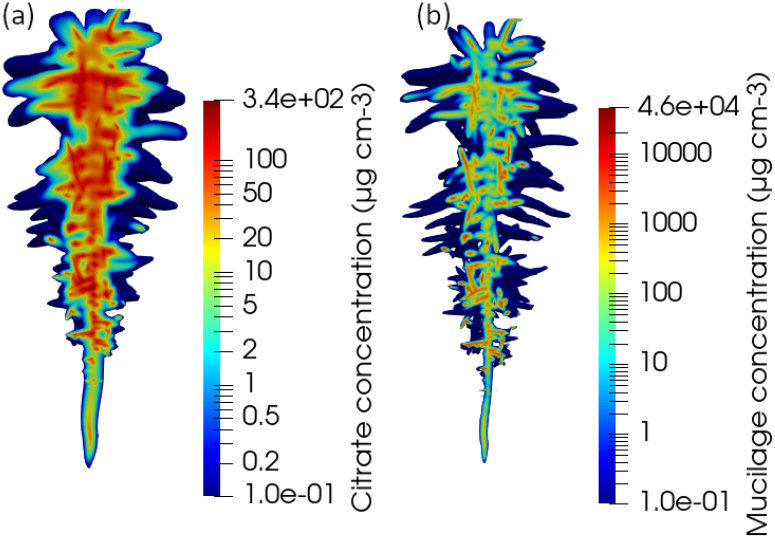
Vertical cut through the distribution of the citrate (a) and mucilage (b) concentrations around the 21 day old root system of *Vicia faba*. Note that the colors are in logarithmic scale and that the color scales differ for the different figures

#### 4.3.0.1 Impact of rhizodeposit overlap on the rhizodeposit hotspot volume

Fig. 6 (a) shows the impact of overlapping rhizodeposition zones on the rhizodeposit hotspot volume of citrate and mucilage around the root system of *Vicia faba*. Interestingly, the impact of overlapping rhizodeposition zones on the total rhizodeposit hotspot volume was much more important for citrate than for mucilage. Furthermore, rhizodeposit hotspot volumes around individual roots were larger for citrate than for mucilage. The relative share of total hotspot volume caused by rhizodeposit overlap increased with increasing simulation time. At simulation day 21, overlapping rhizodeposition zones accounted for 64% of the total citrate rhizodeposit hotspot volume and for 10% of the total mucilage rhizodeposit hotspot volume. Interestingly, the total rhizodeposit hotspot volume without overlap was only slightly higher for citrate than for mucilage. Fig. 6 (b,c) shows the location of overlapping rhizodeposition zones around the root system on the last day of simulation. It can be seen that most of the overlap happened close to the root axis where the branching took place. Rhizodeposit overlap due to individual roots that cross each other freely in the soil domain appeared to be less significant.

**Figure 6:**
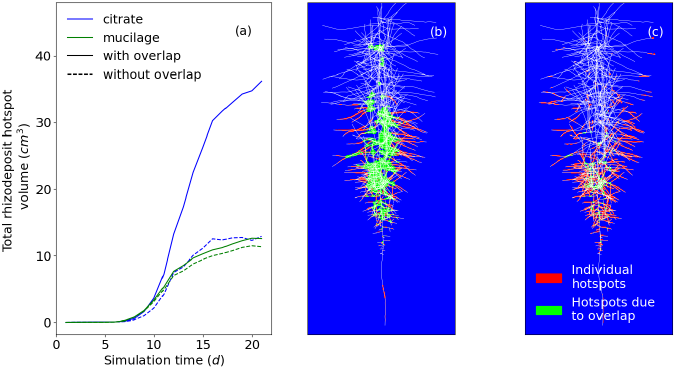
Impact of overlapping rhizodeposition zones on the total rhizodeposit hotspot volume (a), maximal projection along the y-axis of the location of rhizodeposit hotspots caused by overlapping rhizodeposition zones and caused by rhizodeposition from individual roots for citrate (b) and mucilage (c) on simulation day 21

#### 4.3.0.2 Analysis of the duration of rhizodeposit hotspots

The maximum number of days on which hotspot concentrations were reached at a specific location in the soil domain was 16 days for citrate and 9 days for mucilage (Fig. 7 (a)). In general, the longer the duration of the hotspots, the lower was the volume of rhizodeposit hotspots and thus the frequency of rhizodeposit hotspot duration. Interestingly, the most common duration of the rhizodeposit hotspot for mucilage was 3 days. This is the average time between the release of the mucilage at the root tip and its degradation to a concentration below the threshold value. Fig. 7 (b, c) shows the local distribution of the durations of the rhizodeposit hotspots. For both citrate and mucilage, the longest duration of rhizodeposit hotspots occurred near the taproot, where root branching took place and therefore overlapping rhizodeposit zones occurred more frequently. Furthermore, long-lasting rhizodeposit hotspots occurred more frequently around older parts of the root system. Lateral roots of higher order at a greater distance from the taproot did not show long durations of rhizodeposit hotspots. This effect was more pronounced for citrate than for mucilage.

**Figure 7:**
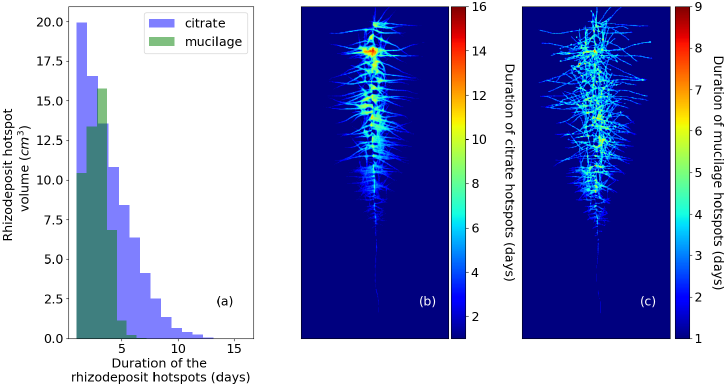
Duration and volume of rhizodeposit hotspots for citrate and mucilage (a); maximal projection along the y-axis of the duration of rhizodeposit hotspots at the different locations in the soil domain for citrate (b) and mucilage (c)

#### 4.3.0.3 Analysis of distance maps from rhizodeposit hotspots

Histograms of distance maps (Fig. 8) of *Vicia faba* show that the volume of soil that was close to a hotspot increased more and more over the simulated 20 day period. At day 5, the small root system and its hotspots were in the top center of the pot. The equidistant surfaces with distances of less than 10 *cm* from the hotspots were approximately semi-spheres around the hotspots, which were at day 5 all near the same point: the parabolic increase of the histogram for less than 10 *cm* distances corresponds to the increase in area of a semi-sphere of radius *r* which is 0.5 × (4*πr*^2^). At a distance of around 10 - 15 *cm*, which corresponds to the phase where the equidistant surface reached the side boundaries of the pot, the histogram line decreases. From 15 - 35 *cm* it remains rather constant and then drops rapidly at a distance of 35 *cm*, which corresponds to the phase where the equidistant surface reached the lower boundary of the pot. At day 10, more and deeper hostspots emerged and as a consequence the peak in the histogram at around 10 *cm* becomes smoother and the drop of the curve occurs now at 25 *cm*. At day 15, the heterogeneous distribution of several hotspots within the domain resulted in a rough histogram line for distances of less than 10 *cm* and hotspots in deeper regions caused a drop at already 15 - 20 *cm* distance where the equidistant surface reached the lower boundary of the pot. Till day 15, the curves for citrate and mucilage were very similar. At day 20, for citrate, there was a peak of the soil volume at a distance of 5 *cm* from the hotspots and for mucilage at a distance of 3 *cm*. At day 20, mucilage showed a larger soil volume in the first five centimeters compared to citrate, which is caused by the wider respectively less clumped distribution of the mucilage hotspots.

**Figure 8:**
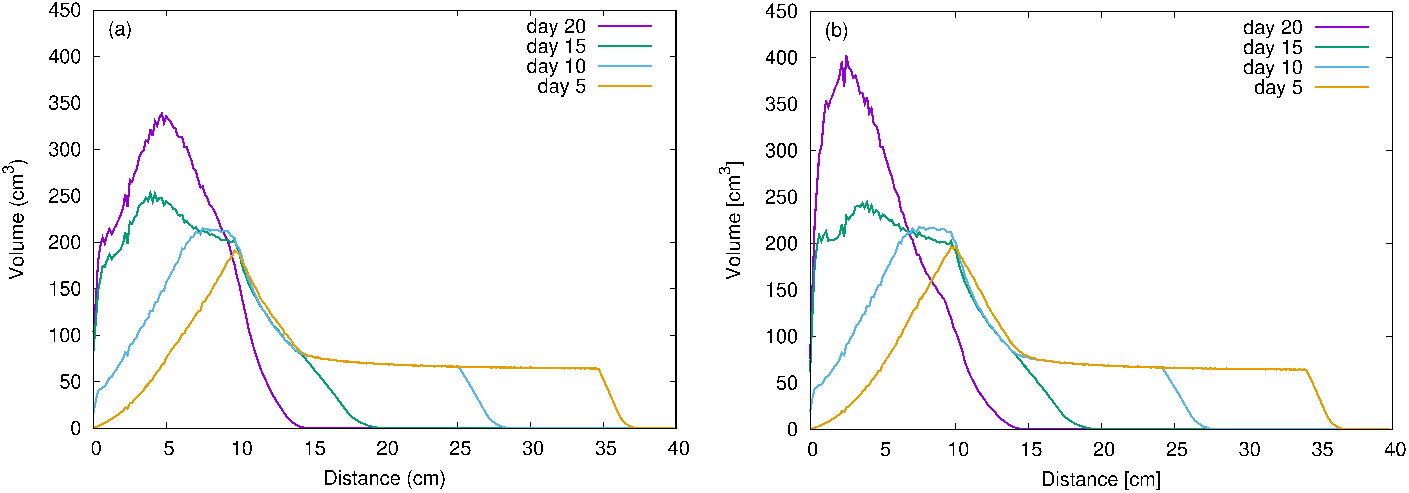
Histograms of distance maps of the Euclidean 3D distance from nearest citrate (a) and mucilage (b) hotspots for *Vicia faba* at day 5, day 10, day 15 and day 20; note that the scales differ in the sub-figures (a) and (b)

## 5. Discussion

### 5.1 The rhizodeposition model

To date, it is not clear how the release of rhizodeposits from an individual root develops with root aging. In our model, we assumed a constant rhizodeposition release rate while the root is growing. As soon as the root stops growing, also rhizodeposition is assumed to stop. Several experimental studies have reported that the total mass of rhizodeposits around a root system is low at the seedling stage of a plant, increases until flowering, and then decreases at maturity (Aulakh et al., 2001; Gransee and Wittenmayer, 2000; Krasil’nikov et al., 1958; Nguyen, 2009). Our model assumptions allow us to simulate such rhizodeposition behaviour and we therefore consider them as justified.

Freshly released mucilage in contact with water is known to diffuse freely into the soil (Sealey et al., 1995). However, when the soil dries, mucilage forms strong bonds between soil particles (Ahmed et al., 2014; Albalasmeh and Ghezzehei, 2014; Sealey et al., 1995). Convective transport of mucilage by flowing water is therefore negligible (Kroener et al., 2018). When microbes decompose mucilage, they are known to simultaneously release gel-like substances called bacterial exopolysaccharides (EPS) (Carminati and Vetterlein, 2013). It has been shown that these substances have similar physical properties to mucilage and are therefore likely to have an effect on the hydraulic properties of the soil (Or et al., 2007). In our study, simulated concentrations of mucilage only refer to fresh mucilage, but not to mucilage derivatives. Similarly, we only considered concentrations of fresh mucilage above the specified threshold value as mucilage hotspots. However, for simulations in which both mucilage deposition and soil water transport are taken into account, the impact of mucilage derivatives on soil hydraulic properties must be considered.

In all simulations, we assumed a constant water content of 0.3 *cm*^3^ *cm*^−3^ in the rhizosphere over the entire simulation period. The assumption of a constant water content is supported by the experimental work of Holz et al. (2018b) and Moradi et al. (2011), who found that the water content in the rhizosphere remained constant regardless of drought stress, which they explained with the high water holding capacity of the mucilage present in the rhizosphere.

In our rhizodeposition model, roots are considered as line sources. The possible influence of the root diameter on the concentration of rhizodeposits is therefore neglected. To satisfy this assumption, the grid resolution used must be larger than the root diameter. On the other hand, a sufficiently fine grid resolution must be chosen to capture the small-scale variations in the spatial distribution of rhizodeposits caused by the steep gradients. Considering that primary roots of *Vicia faba* have a mean root diameter of about 0.95 *mm*, we assumed that a grid resolution of 1 *mm* is suitable to simulate the spatio-temporal distribution of rhizodeposites around the growing root system of *Vicia faba*.

For a soil domain with dimensions of 20 × 20 × 45 *cm*, this resolution resulted in a total number of 1.8 × 10^7^ grid points. For each of these grid points, the rhizodeposit concentration had to be calculated analytically. To keep computation times within acceptable limits, we computed the rhizodeposit concentrations only within a specified radius around each root and parallelized the computation of rhizodeposit concentrations around individual roots.

Our assumption of roots as line sources neglects root diameters and therefore inevitably leads to inaccuracies in the size of the overlap zones of different root types. In addition, the analytical solution is computationally expensive because the rhizodeposition concentrations must be calculated separately for each grid point. To overcome these limitations, the analytical solution could be converted into a numerical approach and integrated into a 3D multicomponent model for solute transport in soil and roots (cf. Mai et al. (2019)). Such a model could then be used to study nutrient uptake by the root system under the influence of dynamic rhizodeposition patterns and, furthermore, to evaluate the influence of differences in root diameter on rhizodeposition patterns.

### 5.2 Rhizodeposition by a single growing root

The differences in the deposition lengths between citrate and mucilage led to differences in the location of the simulated peak concentrations of the two rhizodeposits along the root axis (Table 1, Fig.1 (a)). The maximum simulated radial extent of the mucilage hotspot zone of 0.5 *mm* and the zone where the mucilage concentration was below threshold but still detectable of 2 −5 *mm*, were in the same range as the experimental findings of Holz et al. (2018a) and the calculated values of Zickenrott et al. (2016), which reported rhizosphere extents between 0.6 *mm* and 2 *mm*. For citrate, the maximum simulated radial hotspot extent of 1 *mm* and the detectable concentration extent of 4 − 9 *mm* were of the same order of magnitude as the results for rhizodeposited ^14^*C* from Kuzyakov et al. (2003) who measured a zone of maximum carbon exudate concentration at a distance of 1 − 2 *mm* from the root surface and a zone of less significant amounts of carbon exudate concentration at a distance of 3 − 10 *mm* from the root surface. It must be noted that the experimental conditions and model assumptions in the studies by Holz et al. (2018a), Zickenrott et al. (2016) and Kuzyakov et al. (2003) were not the same as in our modelling setup. They differed with regard to plant species, plant age, water content and pot geometry and may therefore only be regarded as an indicative of the order of magnitude.

### 5.3 Impact of root architectural traits on rhizodeposition patterns

It is well known that root architectural traits have a significant effect on the distribution of rhizodeposits around the root system and thus on rhizosphere processes (Holz et al., 2018b; Lynch, 1995; Nielsen et al., 1994). A detailed analysis about the impact of individual root architectural traits such as root growth rate and branching density on rhizodeposit hotspot volumes and on the rhizodeposition efficiency, however, is still lacking.

Holz et al. (2018b) suggested that reduced root elongation leads to a higher rhizodeposit concentration per rhizosphere soil volume and thus - in the case of mucilage - to an increase in the local water content. In the present study, we made a more detailed analysis of the impact of different root growth rates on the rhizodeposit concentration per rhizosphere soil volume. Considering that a minimum rhizodeposit concentration is required to trigger certain processes, such as an increase in soil water content in the case of mucilage or increased phosphorus mobilization in the case of citrate, an intermediate root growth rate has the greatest effect on rhizosphere processes. If root growth is too fast, the soil volume containing rhizodeposits is large, but the rhizodeposit concentration is below the threshold that triggers a specific rhizosphere process. If root growth is too low, the rhizodeposit concentration is very high, but the soil volume containing such high rhizodeposit concentrations is very low. For our parameterization, the optimal growth rate has been shown to be greater for mucilage than for citrate. It can be speculated that roots take advantage of this effect: When root elongation decreases due to environmental factors, such as soil mechanical impedance, a larger rhizodeposit hotspot volume may result in increased rhizosphere water content in the case of mucilage or increased phosphate availability in the case of citrate, thus compensating for the disadvantages of a smaller root system.

Our simulations showed that an increase in branching density leads to different increases in the total mass of citrate and mucilage in the soil domain. This is due to different release, diffusion, decomposition, and sorption rates of citrate and mucilage. Furthermore, we were able to show that rhizodeposit hotspot volumes around roots that had stopped growing soon disappeared due to the ongoing diffusion and decomposition processes and the resulting decreasing concentrations. In our parameterization, root branching had a greater effect on the total rhizodeposition hotspot volume and also on the rhizodeposition efficiency of citrate than of mucilage. However, if the lateral roots had been shorter, the opposite would have been true because of the difference in deposition length

of citrate and mucilage. Nielsen et al. (1994) and Lynch (1995) reported that highly branched root systems with a large number of root tips have a higher nutrient uptake efficiency and thus a greater influence on rhizosphere processes. Similarly, Fletcher et al. (2020) found that the number of root tips of a root system correlated well with an increase in citrate-enhanced phosphate uptake. This is consistent with the results of our simulations, which also showed larger soil volumes of rhizodeposit hotspots when the number of root tips was increased.

### 5.4 Rhizodeposition patterns around a growing root system

Due to the higher deposition rates (Table 1), the maximum simulated mucilage concentrations were larger than the maximum simulated citrate concentrations which is in line with findings from literature. Zickenrott et al. (2016) estimated that mucilage concentrations of up to 4 × 10^4^ *µg cm*^−3^ soil can potentially occur in the rhizosphere. In our simulations, the maximum observed mucilage concentration amounted to 2.7 × 10^5^ *µg cm*^−3^ soil and is therefore a bit higher than this estimated maximum value. Gerke (2015) and Jones (1998) found maximum citrate concentrations in the rhizosphere between 1 × 10^3^ and 4 × 10^3^ *µg cm*^−3^ soil. These ranges are a bit higher than our maximum simulated citrate concentration of 938 *µg cm*^−3^ soil. This can be explained by the fact that other plants such as *Lupinus albus* and *Cicer arietinum* have been shown to release much greater amounts of citrate into the soil than *Vicia faba*.

The rhizodeposit hotspot analysis showed the importance of overlapping rhizodeposition zones for the development of rhizodeposit hotspots. The overlap of rhizodeposits was shown to account for 64 % of the total volume of rhizodeposits of citrate, but only for 10 % of the total volume of rhizodeposits of mucilage after 21 simulation days. This difference is caused primarily by differences in the rhizodeposit release: while mucilage is deposited exclusively at the root tip, citrate release takes place over a length of approximately 5 *cm* behind the root tip. Additionally, due to the larger diffusion coefficient of citrate compared to mucilage, rhizodeposit concentration volumes around individual roots are larger for citrate than for mucilage and the possibility of rhizodeposit overlap is thus also greater for citrate than for mucilage. In the case of high branching densities, it can be assumed that individual hotspot volumes around roots will overlap, thereby leading to a decrease in the total rhizodeposit hotspot volume. For our parameterization, however, the hotspot volumes that were created by rhizodeposition overlap were more important than the hotspot volumes that were lost by rhizodeposition overlap. Due to the increasing number of laterals, the relative share of total hotspot volume caused by rhizodeposit overlap was shown to increase with increasing simulation time for our parameterization. It must be noted that we only looked at a single root system in the present study. If multiple neighbouring root systems were considered, the impact of overlapping rhizodeposition zones on the total rhizodeposit hotspot volume would be even larger. Our simulations have shown that long-lasting rhizodeposit hotspots occur mainly in that part of the root system where branching occurs and where overlapping rhizodeposition zones are therefore more frequent. In our example root system *Vicia faba*, the zone of long-lasting rhizodeposit hotspots is thus found near the taproot, where lateral roots emerge. It can therefore be expected that rhizosphere processes such as an increase in soil water content in the case of mucilage or increased phosphorus mobilization in the case of citrate are stronger within the part of a root system where branching takes place. The analysis of distance maps of rhizodeposit hotspots showed that the characteristics of a specific rhizodeposit have a significant effect on the distribution of distances from any point in the soil domain to the nearest rhizodeposit hotspot: Mucilage hotspots were found to be more widely distributed in the soil domain than citrate hotspots, and therefore had a larger soil volume with a short distance to the nearest hotspot. Considering that certain bacteria in soil can respond to organic compounds detected from a certain distance, these results are significant for microbially controlled processes in the rhizosphere.

There are numerous modeling studies in the literature on root foraging strategies that use 3D root architecture models (e.g. Ge et al. (2000), Lynch (1995), and Pagés (2011)). However, all of these studies concentrated on the analysis of nutrient depletion zone overlap and did not consider the impact of overlapping rhizodeposition zones on nutrient supply. De Parseval et al. (2017) used a 2D model approach to investigate the interaction between inter-root competition and inter-root facilitation in the horizontal plane. Inter-root competition is caused by the overlap of nutrient depletion zones, while inter-root facilitation is based on the overlap of rhizodeposition zones, which leads to rhizodeposit hotspots and consequently to an increased nutrient availability. Based on the distances between roots, this model approach allowed them to predict whether competition, facilitation or no interaction is the predominant process governing root phosphorus uptake. It would be pertinent to use our model to bridge these studies and to extend previous modelling approaches on root for-aging strategies by the aspect of inter-root facilitation. This would give us a more realistic estimate about the impact of root architecture on root nutrient uptake.

## 5.5 Conclusion

In this study, we presented a new model to simulate the spatiotemporal distribution patterns of rhizodeposits around growing root systems in three dimensions. The novel model approach allowed us to evaluate the effects of root architecture features such as root growth rate and branching density on the development of rhizodeposit hotspot zones, which can trigger specific rhizosphere processes such as increased nutrient uptake by roots. It further enables the investigation of the influence of differences in rhizodeposit properties and root architectures of different plant species on rhizodeposition patterns. We showed that rhizodeposit hotspot volumes around roots were at a maximum at intermediate root growth rates and that branching allowed the rhizospheres of individual roots to overlap, resulting in an increase in the volume of rhizodeposit hotspot zones.

In the future we aim to intergate our model into a 3D multi-component root and solute transport model (Mai et al., 2019). This model could then be used to mechanistically explain experimentally observed rhizodeposition patterns (e.g., using zymography or ^11^*CO*2-labeling (Giles et al., 2018; Yin et al., 2020)). We also aim to incorporate the influence of root hairs and root diameters into our model to gain a better understanding of the water and nutrient acquisition strategies of different plant species.

## 5.6 Acknowledgements

This project was carried out in the framework of the priority program 2089 ‘Rhizosphere spatiotemporal organization – a key to rhizosphere functions’ funded by the German Research Foundation DFG under the project number 403641034. A.H. and E.K. have been funded by the German Research Foundation DFG under Germany’s Excellence Strategy – EXC 2070 – 390732324.

## References

Ahmed, M. A., Kroener, E., Holz, M., Zarebanadkouki, M., & Carminati, A. (2014). Mucilage exudation facilitates root water uptake in dry soils. Functional Plant Biology, 41 (11), 1129–1137. https://doi.org/10.1071/FP13330

Ahmed, M. A., Kroener, E., Benard, P., Zarebanadkouki, M., Kaestner, A., & Carminati, A. (2016). Drying of mucilage causes water repellency in the rhizosphere of maize: Measurements and modelling. Plant and Soil, 407 (1-2), 161–171. https://doi.org/10.1007/s11104-015-2749-1

Albalasmeh, A. A., & Ghezzehei, T. A. (2014). Interplay between soil drying and root exudation in rhizosheath development. Plant and Soil, 374 (1-2), 739–751. https://doi.org/10.1007/s11104-013-1910-y

Aulakh, M., Wassmann, R., Bueno, C., Kreuzwieser, J., & Rennenberg, H. (2001). Characterization of root exudates at different growth stages of ten rice (oryza sativa l.) cultivars. Plant biology, 3 (2), 139–148. https://doi.org/10.1055/s-2001-12905:

Bear, J., & Cheng, A. H.-D. (2010). Modeling groundwater flow and contaminant transport (Vol. 23). Springer Science & Business Media.

Benard, P., Zarebanadkouki, M., Brax, M., Kaltenbach, R., Jerjen, I., Marone, F., Couradeau, E., Felde, V. J., Kaestner, A., & Carminati, A. (2019). Microhydrological niches in soils: How mucilage and eps alter the biophysical properties of the rhizosphere and other biological hotspots. Vadose Zone Journal, 18 (1), 1–10. https://doi.org/10.2136/vzj2018.12.0211

Carminati, A., Kroener, E., Ahmed, M. A., Zarebanadkouki, M., Holz, M., & Ghezzehei, T. (2016). Water for carbon, carbon for water. Vadose Zone Journal, 15 (2), 1–10. https://doi.org/10.2136/vzj2015.04.0060

Carminati, A., & Vetterlein, D. (2013). Plasticity of rhizosphere hydraulic properties as a key for efficient utilization of scarce resources. Annals of botany, 112 (2), 277–290. https://doi.org/10.1093/aob/mcs262

Carslaw, H., & Jaeger, J. (1959). Conduction of heat in solids. Clarendon Press. https://doi.org/10.1007/978-3-319-48090-99

Carson, J. K., Gonzalez-Quiñones, V., Murphy, D. V., Hinz, C., Shaw, J. A., & Gleeson, D. B. (2010). Low pore connectivity increases bacterial diversity in soil. Applied and environmental microbiology, 76 (12), 3936–3942. https://doi.org/10.1128/AEM.03085-09

Cheng, W., & Gershenson, A. (2007). Carbon fluxes in the rhizosphere. The rhizosphere (pp. 31–56). Elsevier. https://doi.org/0.1016/B978-012088775-0/50004-5

Crank, J. (1979). The mathematics of diffusion. Oxford university press.

Darrah, P. (1991). Models of the rhizosphere. Plant and Soil, 133 (2), 187–199.

De Parseval, H., Barot, S., Gignoux, J., Lata, J.-C., & Raynaud, X. (2017). Modelling facilitation or competition within a root system: Importance of the overlap of root depletion and accumulation zones. Plant and soil, 419 (1-2), 97–111. https://doi.org/10.1007/s11104-017-3321-y

Evans, L. C. (1998). Partial differential equations. Graduate studies in mathematics, 19 (2).

Fletcher, D. M., Ruiz, S., Dias, T., Petroselli, C., & Roose, T. (2020). Linking root structure to functionality: The impact of root system architecture on citrate-enhanced phosphate uptake. New Phytologist, 227 (2), 376–391. https://doi.org/doi.org/10.1111/nph.16554

Fletcher, D. M., Shaw, R., Sánchez-Rodriguez, A., Daly, K., Van Veelen, A., Jones, D., & Roose, T. (2019). Quantifying citrate-enhanced phosphate root uptake using microdialysis. Plant and Soil, 1–21. https://doi.org/10.1007/s11104-019-04376-4

Gao, W., Blaser, S. R., Schlüter, S., Shen, J., & Vetterlein, D. (2019). Effect of localised phosphorus application on root growth and soil nutrient dynamics in situ–comparison of maize (zea mays) and faba bean (vicia faba) at the seedling stage. Plant and Soil, 441 (1-2), 469–483. [https](https://doi.org/10.1007/s11104-019-04138-2)

Ge, Z., Rubio, G., & Lynch, J. P. (2000). The importance of root gravitropism for inter-root competition and phosphorus acquisition efficiency: Results from a geometric simulation model. Plant and soil, 218 (1-2), 159–171. https://doi.org/10.1023/A:1014987710937

Gerke, J. (2015). The acquisition of phosphate by higher plants: Effect of carboxylate release by the roots. a critical review. Journal of Plant Nutrition and Soil Science, 178 (3), 351–364. https://doi.org/10.1002/jpln201400590.

Giles, C., Dupuy, L., Boitt, G., Brown, L., Condron, L., Darch, T., Blackwell, M., Menezes-Blackburn, D., Shand, C., Stutter, M., et al. (2018). Root development impacts on the distribution of phosphatase activity: Improvements in quantification using soil zymography. Soil Biology and Biochemistry, 116, 158–166. https://doi.org/10.1016/j.soilbio.2017.08.011

Gransee, A., & Wittenmayer, L. (2000). Qualitative and quantitative analysis of water-soluble root exudates in relation to plant species and development. Journal of plant nutrition and soil science, 163 (4), 381–385. https://doi.org/10.1002/1522-2624(200008)163:4381::AID-JPLN3813.0.CO;2-7

Hinsinger, P., Bengough, A. G., Vetterlein, D., & Young, I. M. (2009). Rhizosphere: Biophysics, biogeochemistry and ecological relevance. Plant and soil, 321 (1-2), 117–152. https://doi.org/10.1007/s11104-008-9885-9

Hodge, A., Berta, G., Doussan, C., Merchan, F., & Crespi, M. (2009). Plant root growth, architecture and function. Plant and soil, 321 (1-2), 153–187. https://doi.org/10.1007/s11104-009-9929-9

Holz, M., Leue, M., Ahmed, M. A., Benard, P., Gerke, H. H., & Carminati, A. (2018a). Spatial distribution of mucilage in the rhizosphere measured with infrared spectroscopy. Frontiers in Environmental Science, 6, 87. https://doi.org/10.3389/fenvs.2018.00087

Holz, M., Zarebanadkouki, M., Kaestner, A., Kuzyakov, Y., & Carminati, A. (2018b). Rhizodeposition under drought is controlled by root growth rate and rhizosphere water content. Plant and soil, 423 (1-2), 429–442. https://doi.org/10.1007/s11104-017-3522-4

Iijima, M., Sako, Y., & Rao, T. (2003). A new approach for the quantification of root-cap mucilage exudation in the soil. Roots: The dynamic interface between plants and the earth (pp. 399–407). Springer. https://doi.org/10.1023/A:1026183109329

Jacques, D., Šimnek, J., Mallants, D., & Van Genuchten, M. T. (2018). The hpx software for multicomponent reactive transport during variably-saturated flow: Recent developments and applications. Journal of Hydrology and Hydromechanics, 66 (2), 211–226. https://doi.org/10.1515/johh-2017-0049

Jones, D. L. (1998). Organic acids in the rhizosphere–a critical review. Plant and soil, 205 (1), 25–44. https://doi.org/10.1023/A:1004356007312

Kirk, G. D. (1999). A model of phosphate solubilization by organic anion excretion from plant roots. European Journal of Soil Science, 50 (3), 369–378. https://doi.org/10.1111/j.1365-2389.1999.00239.x

Krasil’nikov, N. et al.. (1958). Soil microorganisms and higher plants. Soil microorganisms and higher plants.

Kroener, E., Holz, M., Zarebanadkouki, M., Ahmed, M., & Carminati, A. (2018). Effects of mucilage on rhizosphere hydraulic functions depend on soil particle size. Vadose Zone Journal, 17 (1), 1–11. https://doi.org/102136/vzj2017.03.0056.

Kuzyakov, Y., Raskatov, A., & Kaupenjohann, M. (2003). Turnover and distribution of root exudates of zea mays. Plant and Soil, 254 (2), 317–327. https://doi.org/10.1023/A:1025515708093

Lobet, G., Pound, M. P., Diener, J., Pradal, C., Draye, X., Godin, C., Javaux, M., Leitner, D., Meunier, F., Nacry, P., et al. (2015). Root system markup language: Toward a unified root architecture description language. Plant Physiology, 167 (3), 617–627. https://doi.org/10.1104/pp114.253625.

Lynch, J. (1995). Root architecture and plant productivity. Plant physiology, 109 (1), 7. https://doi.org/10.1104/pp.109.1.7

Lynch, J. P., Ho, M. D. et al.. (2005). Rhizoeconomics: Carbon costs of phosphorus acquisition. Plant and soil, 269 (1-2), 45–56. https://doi.org/10.1007/s11104-004-1096-4

Lyu, Y., Tang, H., Li, H., Zhang, F., Rengel, Z., Whalley, W. R., & Shen, J. (2016). Major crop species show differential balance between root morphological and physiological responses to variable phosphorus supply. Frontiers in Plant Science, 7, 1939. https://doi.org/10.3389/fpls.201601939.

Mai, T. H., Schnepf, A., Vereecken, H., & Vanderborght, J. (2019). Continuum multiscale model of root water and nutrient uptake from soil with explicit consideration of the 3d root architecture and the rhizosphere gradients. Plant and Soil, 439 (1-2), 273–292. https://doi.org/10.1007/s11104-018-3890-4

Manschadi, A. M., Kaul, H.-P., Vollmann, J., Eitzinger, J., & Wenzel, W. (2014). Developing phosphorus-efficient crop varieties—an interdisciplinary research framework. Field Crops Research, 162, 87–98. https://doi.org/10.1016/j.fcr.2013.12.016

Moradi, A. B., Carminati, A., Vetterlein, D., Vontobel, P., Lehmann, E., Weller, U., Hopmans, J. W., Vogel, H.-J., & Oswald, S. E. (2011). Threedimensional visualization and quantification of water content in the rhizosphere. New Phytologist, 192 (3), 653–663. https://doi.org/10.1111/j.1469-8137.2011.03826.x

Nguyen, C. (2009). Rhizodeposition of organic c by plant: Mechanisms and controls. Sustainable agriculture (pp. 97–123). Springer. https://doi.org/10.1007/978-90-481-2666-89.

Nguyen, C., Froux, F., Recous, S., Morvan, T., & Robin, C. (2008). Net n immobilisation during the biodegradation of mucilage in soil as affected by repeated mineral and organic fertilisation. Nutrient Cycling in Agroecosystems, 80 (1), 39–47. https://doi.org/10.1007/s10705-007-9119-1

Nielsen, K. L., Lynch, J. P., Jablokow, A. G., & Curtis, P. S. (1994). Carbon cost of root systems: An architectural approach. Plant and Soil, 165 (1), 161–169. https://doi.org/10.1007/BF00009972

Nye, P. (1966). The effect of the nutrient intensity and buffering power of a soil, and the absorbing power, size and root hairs of a root, on nutrient absorption by diffusion. Plant and Soil, 25 (1), 81–105. https://doi.org/10.1007/BF01347964

Oburger, E., Jones, D. L., & Wenzel, W. W. (2011). Phosphorus saturation and ph differentially regulate the efficiency of organic acid anion-mediated p solubilization mechanisms in soil. Plant and Soil, 341 (1-2), 363–382. https://doi.org/10.1007/s11104-010-0650-5

Olesen, T., Moldrup, P., Yamaguchi, T., & Rolston, D. (2001). Constant slope impedance factor model for predicting the solute diffusion coefficient in unsaturated soil. Soil Science, 166 (2), 89–96. https://doi.org/10.1097/00010694-200102000-00002

Ollion, J., Cochennec, J., Loll, F., Escudé, C., & Boudier, T. (2013). Tango: A generic tool for high-throughput 3d image analysis for studying nuclear organization. Bioinformatics, 29 (14), 1840–1841. https://doi.org/10.1093/bioinformatics/btt276

Or, D., Phutane, S., & Dechesne, A. (2007). Extracellular polymeric substances affecting pore-scale hydrologic conditions for bacterial activity in unsaturated soils. Vadose Zone Journal, 6 (2), 298–305. https://doi.org/10.2136/vzj2006.0080

Pagès, L. (2011). Links between root developmental traits and foraging performance. Plant, Cell & Environment, 34 (10), 1749–1760.

Pausch, J., Tian, J., Riederer, M., & Kuzyakov, Y. (2013). Estimation of rhizodeposition at field scale: Upscaling of a 14 c labeling study. Plant and Soil, 364 (1-2), 273–285. https://doi.org/10.1007/s11104-012-1363-8

Pineros, M. A., Magalhaes, J. V., Alves, V. M. C., & Kochian, L. V. (2002). The physiology and biophysics of an aluminum tolerance mechanism based on root citrate exudation in maize. Plant Physiology, 129 (3), 1194–1206. https://doi.org/10.1104/pp.002295

Rangel, A. F., Rao, I. M., Braun, H.-P., & Horst, W. J. (2010). Aluminum resistance in common bean (phaseolus vulgaris) involves induction and maintenance of citrate exudation from root apices. Physiologia Plantarum, 138 (2), 176–190. https://doi.org/10.1111/j.1399-3054.2009.01303.x

Schindelin, J., Arganda-Carreras, I., Frise, E., Kaynig, V., Longair, M., Pietzsch, T., Preibisch, S., Rueden, C., Saalfeld, S., Schmid, B., et al. (2012). Fiji: An open-source platform for biological-image analysis. Nature methods, 9 (7), 676–682. https://doi.org/10.1038/nmeth.2019

Schnepf, A., Leitner, D., & Klepsch, S. (2012). Modeling phosphorus uptake by a growing and exuding root system. Vadose Zone Journal, 11 (3), vzj2012–0001. https://doi.org/10.2136/vzj2012.0001

Schnepf, A., Leitner, D., Landl, M., Lobet, G., Mai, T. H., Morandage, S., Sheng, C., Zörner, M., Vanderborght, J., & Vereecken, H. (2018). Crootbox: A structural–functional modelling framework for root systems. Annals of botany, 121 (5), 1033–1053. https://doi.org/10.1093/aob/mcx221

Schulz-Bohm, K., Geisen, S., Wubs, E. J., Song, C., de Boer, W., & Garbeva, P. (2017). The prey’s scent–volatile organic compound mediated interactions between soil bacteria and their protist predators. The ISME journal, 11 (3), 817–820. https://doi.org/10.1038/ismej.2016.144

Sealey, L., McCully, M., & Canny, M. (1995). The expansion of maize root-cap mucilage during hydration. 1. kinetics. Physiologia Plantarum, 93 (1), 38–46. https://doi.org/10.1034/j.1399-3054.1995.930107.x

Stingaciu, L., Schulz, H., Pohlmeier, A., Behnke, S., Zilken, H., Javaux, M., & Vereecken, H. (2013). In situ root system architecture extraction from magnetic resonance imaging for water uptake modeling. Vadose zone journal, 12 (1), 1–9. https://doi.org/10.2136/vzj2012.0019

Watt, M., Silk, W. K., & Passioura, J. B. (2006). Rates of root and organism growth, soil conditions, and temporal and spatial development of the rhizosphere. Annals of botany, 97 (5), 839–855. https://doi.org/10.1093/aob/mcl028

Westhoff, S., van Wezel, G. P., & Rozen, D. E. (2017). Distance-dependent danger responses in bacteria. Current opinion in microbiology, 36, 95–101. https://doi.org/10.1016/j.mib.2017.02.002

Wilson, J. L., & Miller, P. J. (1978). Two-dimensional plume in uniform groundwater flow. Journal of the Hydraulics Division, 104 (4), 503–514. [https](https://doi.org/10.1061/JYCEAJ.0004975)

Yin, Y.-G., Suzui, N., Kurita, K., Miyoshi, Y., Unno, Y., Fujimaki, S., Nakamura, T., Shinano, T., & Kawachi, N. (2020). Visualising spatio-temporal distributions of assimilated carbon translocation and release in root systems of leguminous plants. Scientific reports, 10 (1), 1–11. https://doi.org/10.1038/s41598-020-65668-9:

Zhou, X.-R., Schnepf, A., Vanderborght, J., Leitner, D., Lacointe, A., Vereecken, H., & Lobet, G. (2020). Cplantbox, a whole-plant modelling framework for the simulation of water-and carbon-related processes. in silico Plants, 2 (1), diaa001. https://doi.org/10.1093/insilicoplants/diaa001

Zickenrott, I.-M., Woche, S. K., Bachmann, J., Ahmed, M. A., & Vetterlein, D. (2016). An efficient method for the collection of root mucilage from different plant species—a case study on the effect of mucilage on soil water repellency. Journal of Plant Nutrition and Soil Science, 179 (2), 294–302. https://doi.org/10.1002/jpln.201500511

