## Supplementary Material for "Simulating rhizodeposition patterns around growing and exuding root systems"

Table S1: CPlantBox root growth parameters for *Vicia faba* and *Zea mays*

| Description | Symbol | Unit | Vicia faba | Zea mays |
|---|---|---|---|---|
| Time delay between the basal roots | *delayB* | *day* | – | 0.786 |
| Emergence of first basal root | *firstB* | *day* | – | 0.108 |
| Maximal number of basal roots | *maxB* | - | – | 15.75 |
| | | | **taproot** | **primary / basal roots** |
| Basal zone | *lb* | *cm* | 1.00 (0.50) | 1.38 (1.24) |
| Apical zone | *la* | *cm* | 9.97 (1.99) | 2.23 (1.21) |
| Internodal distance | *ln* | *cm* | 0.16 (0.22) | 0.57 (0.20) |
| Root radius | *a* | *cm* | 0.04 (0.02) | 0.04 (0.02) |
| Branching angle | $\theta$ | *rad* | 0 (0) | 1.20 (0.10) |
| Growth function | *gf* | - | 1 | 2 |
| Initial growth rate | *r* | $cmd^{-1}$ | 2.07 (0) | 1.60 (0) |
| Maximal root length | *lmax* | *cm* | 150 (0) | 150 (0) |
| Type of tropism | *tropismT* | - | 1 | 1 |
| Number of trials | *tropismN* | - | 1.8 | 1.5 |
| Mean value of expected tropism change | *tropismS* | $cm^{-1}$ | 0.2 | 0.2 |
| | | | **1st order laterals** | **1st order laterals** |
| Basal zone | *lb* | *cm* | 1 (0) | 1.25 (0) |
| Apical zone | *la* | *cm* | 2 (0) | 1.04 (0) |
| Internodal distance | *ln* | *cm* | 5 (0) | 0.65 (0.87) |
| Root radius | *a* | *cm* | 0.05 (0.006) | 0.03 (0.01) |
| Branching angle | $\theta$ | *rad* | 1.42 (0) | 1.47 (0.55) |
| Growth function | *gf* | - | 1 | 2 |
| Initial growth rate | *r* | $cmd^{-1}$ | 0.82(0.08) | 1.88 (0.06) |
| Maximal root length | *lmax* | *cm* | 8 (0.8) | 2.29 (1.81) |
| Type of tropism | *tropismT* | - | 2 | 2 |
| Number of trials | *tropismN* | - | 0.5 | 2 |
| Mean value of expected tropism change | *tropismS* | $cm^{-1}$ | 0.2 | 0.69 |
| | | | **2nd order laterals** | **2nd order laterals** |
| Basal zone | *lb* | *cm* | 0 (0) | 0 (0) |
| Apical zone | *la* | *cm* | 10 (0) | 10 (0) |
| Internodal distance | *ln* | *cm* | 100 | 100 |
| Root radius | *a* | *cm* | 0.03 (0) | 0.03 (0) |
| Branching angle | $\theta$ | *rad* | 1.42 (0) | 1.50 (0.15) |
| Growth function | *gf* | - | 1 | 2 |
| Initial growth rate | *r* | $cmd^{-1}$ | 0.82(0.08) | 2.35 (0.08) |
| Maximal root length | *lmax* | *cm* | 4 (0.4) | 0.94 (0) |
| Type of tropism | *tropismT* | - | 2 | 2 |
| Number of trials | *tropismN* | - | 0.5 | 0.5 |
| Mean value of expected tropism change | *tropismS* | $cm^{-1}$ | 0.2 | 0.1 |

### S1 Auxiliary study: Comparison of citrate and mucilage rhizodeposition by the tap root system of *Vicia faba* and the fibrous root system of *Zea mays*

In analogy to *Vicia faba*, root architecture parameters were obtained from µCT images from the lab experiment by Gao et al. (2019). Rhizodeposit release rates of *Zea mays* were set to $3.7\mu g\, d^{-1}\, cm\, root^{-1}$ for citrate (Pineros et al., 2002) and to $5.27\mu g\, d^{-1}\, root\, tip^{-1}$ for mucilage (Zickenrott et al., 2016). All other model input parameters were the same as for the simulations with *Vicia faba*.

Fig. S1 shows the rhizodeposit concentration patterns of citrate and mucilage around the 21 day old root system of *Zea mays*. The maximum extent of the rhizosphere was defined using an arbitrary threshold of $0.1\, \mu g\, cm^{-3}$.

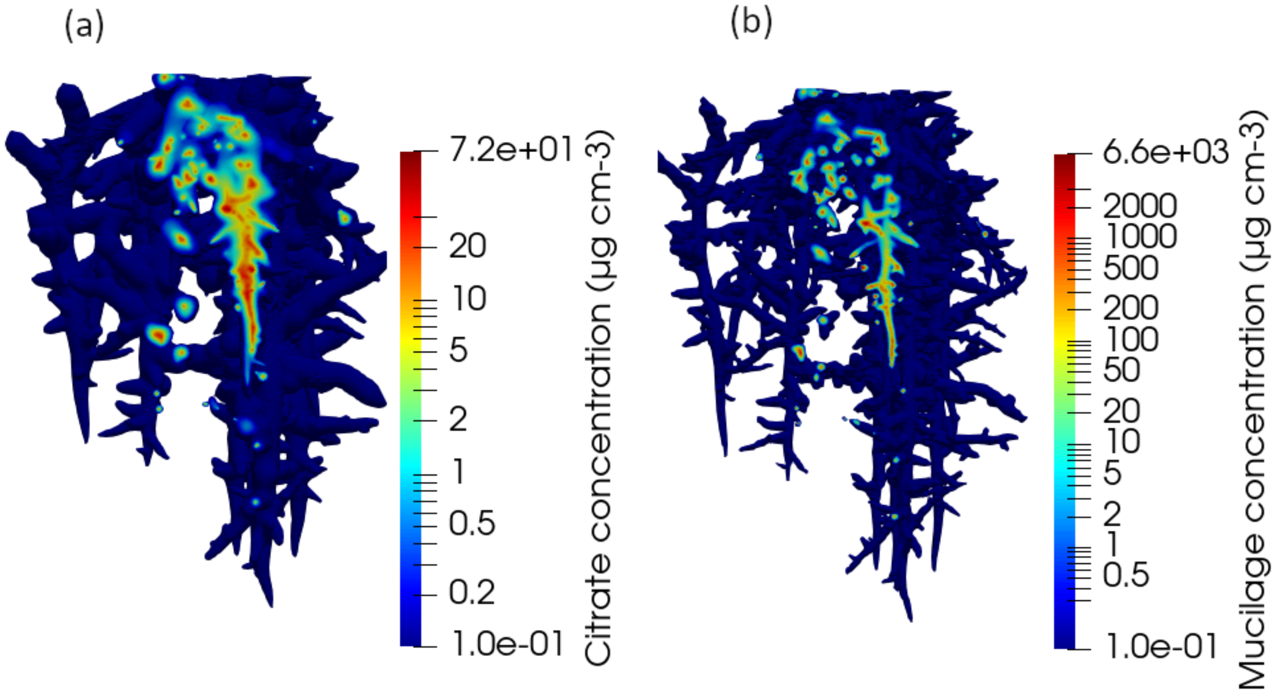


Figure S1: Vertical cut through the distribution of the citrate (a) and mucilage (b) concentrations around the 21 day old root system of *Zea mays*. Note that the colors are in logarithmic scale and that the color scales differ for the different figures

Fig. S2 shows the amount of released citrate and mucilage rhizodeposits from the root systems of *Vicia faba* and *Zea mays* with time. The total mass of rhizodeposits present in the soil domain gradually increases while the root system is growing. It is larger for mucilage than for citrate and mostly also larger for *Vicia faba* than for *Zea mays*. Only between simulation day 5 and simulation day 8, the emergence time of lateral roots of *Vicia faba*, the total rhizodeposit mass is larger for *Zea mays* than for *Vicia faba* (Fig. S2 (a)). The total mass of rhizodeposits normalized with the total root length shows very distinct patterns for *Vicia faba* and *Zea mays* (Fig. S2 (b)). For *Vicia faba*, the curve clearly reflects the development of the root architecture: At simulation day 6, the first lateral roots emerge, which is reflected by a sharp increase in the root length-normalized mucilage mass. For citrate, which is released over a length of $5\, cm$ behind the root apex, this increase can be seen to a lesser extent and with a certain delay. For *Zea mays*, the length-normalized citrate and mucilage masses remain relatively constant over the entire simulation period, which is caused by the large number of basal roots and the early emergence of first order laterals at simulation day 3, which level out any visible impact of root architecture. Similar patterns arise for the total mass of rhizodeposits normalized with the number of root tips (Fig. S2 (c)). For *Vicia faba*, the emergence of first and second order lateral roots (simulation day 6 and 7, respectively), is reflected in the curves of both citrate and mucilage. For *Zea mays*, the curves are relatively stable over the entire simulation period. Fig. S2 (d) shows the total mass of rhizodeposits normalized with the volume of the convex hull of the root system. At the beginning of the simulation period, the values are extremely large due to the small volume of the convex hull, but they level out at approximately simulation day

7. It can be seen that for both citrate and mucilage, the convex hull normalized rhizodeposit mass and thus the rhizodeposit concentrations are larger for *Vicia faba* than for *Zea mays*.

On simulation day 21, the root system of *Zea mays* was 2 times longer, had 3.5 times more root tips, and had a convex hull volume 3.7 times larger than the root system of *Vicia faba* (Fig. S2 (b,c,d), red curves). However, the total mass of released citrate and mucilage was only 11 % and 14 % of that of *Vicia faba*, respectively (Fig. S2 (a)). According to our simulations, the larger root system of *Zea mays* could therefore not make up for the lower rhizodeposit release rate as compared to *Vicia faba* to reach similar amounts of rhizodeposit mass released into the soil.

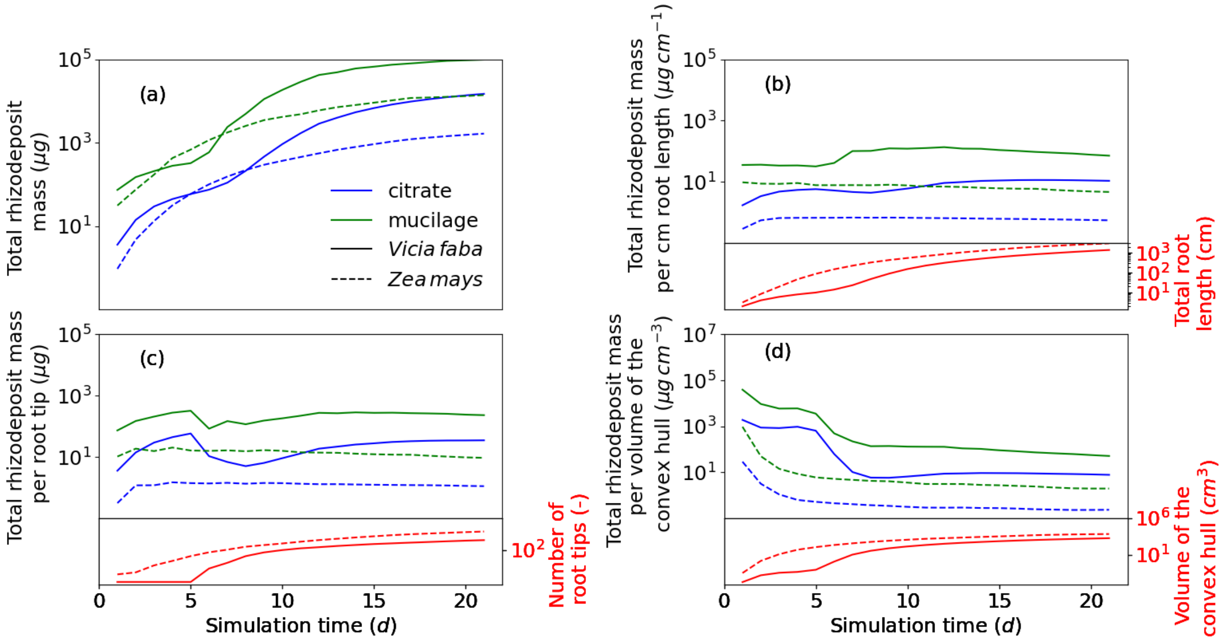


Figure S2: Total amount of released rhizodeposit mass over time (a), normalized with the total root length (b), normalized with the number of root tips (c) and normalized with the volume of the convex hull (d) for citrate and mucilage the root systems *Vicia faba* and *Zea mays*; note that all y-axes are in logarithmic scale
